# Whole genome resequencing of a laboratory-adapted *Drosophila melanogaster* population sample

**DOI:** 10.1101/081554

**Authors:** William P. Gilks, Tanya M. Pennell, Ilona Flis, Matthew T. Webster, Edward H. Morrow

## Abstract

As part of a study into the molecular genetics of sexually dimorphic complex traits, we used next-generation sequencing to obtain data on genomic variation in an outbred laboratory-adapted fruit fly (*Drosophila melanogaster*) population. We successfully resequenced the whole genome of 2 females from the Berkeley reference line (BDGP6/dm6), and 220 hemiclonal females that were heterozygous for the same reference line genome, and a unique haplotype from the outbred base population (LH_M_). The use of a static and known genetic background enabled us to obtain sequences from whole-genome phased haplotypes. We used a BWA-Picard-GATK pipeline for mapping sequence reads to the dm6 reference genome assembly, at a median depth-of coverage of 31X, and have made the resulting data publicly-available in the NCBI Short Read Archive (BioProject PRJNA282591). Haplotype Caller discovered and genotyped 1,726,931 genetic variants (SNPs and indels, <200bp). Additionally, we used GenomeStrip/2.0 to discover and genotype 167 large structural variants (1-100Kb in size). Sequence data and quality-filtered genotype data are publicly-available at NCBI (Short Read Archive, dbSNP and dbVar). We have also released the unfiltered genotype data, and the code and logs for data processing, summary statistics, and graphs, via the research data repository, Zenodo, (https://zenodo.org/, ’Sussex Drosophila Sequencing’ community).

## 1 Introduction

As part of a study on the molecular genetics of sexually dimorphic complex traits, we used hemiclonal analysis in conjunction with next-generation sequencing to characterise molecular genetic variation across the genome, from an outbred laboratory-adapted population of *Drosophila melanogaster,* known as LH_M_^1,11^. The hemiclone experimental design allows the repeated phenotyping of multiple individuals, each with the same unrecombined haplotype on a different random genetic background. This method has been used to investigate standing genetic variation and intersexual genetic correlations for quantitative traits^1^ and gene expression^7^, but it has not yet been used to obtain genomic data.

The 220 hemiclone females that were sequenced in the present study have a maternal haplotype, from the *dm6* reference assembly strain (BDGP6+ISO1 mito/dm6, Bloomington *Drosophila* Stock Center no. 2057)^2,6^, and have a different paternal genome each, sampled using cytogenetic cloning from the LH_M_ base population. All non-reference genotypes in the sequenced LH_M_ hemiclones were expected to be heterozygous and in-phase, except in rare instances where the in-house dm6 reference strain also had the same non-reference allele.

Previous studies indicate that the limits for DNA quantity in 'next-generation' sequencing are 50-500ng^12^. We sequenced individual *D.melanogaster,* rather than pools of clones, because more biological information can be obtained, and because modern transposon-based library preparation allows accurate sequencing at low concentrations of DNA. *D. melanogaster* is a small insect (~1μg) although this problem is off-set by the reduced proportion of repetitive intergenic sequence, and small genome size relative to other insects (170Mb verses ~500Mb),^12^.

We mapped reads to the *D. melanogaster* dm6 reference assembly using a BWA-Picard-GATK pipeline, and called nucleotide variants using both HaplotypeCaller, and Genomestrip, the latter of which detections copy-number variation up to 1Mb in length. We have made the mapped sequencing data, and genotype data publicly-available on NCBI, and additionally have made the meta-data, analysis code and logs publicly-available on the research data repository, Zenodo. This is the first report of a study which uses methods for detecting both SNPs, indels and CNVs genome-wide in next-generation sequencing data, and the first report of whole genome resequencing in hemiclonal individuals.

## 2 Materials and Methods

### 2.1 Study samples

The base population (LH_M_) was originally established from a set of 400 inseminated females, trapped by Larry Harshman in a citrus orchard near Escalon, California in 1991^11^. It was initially kept at a large size (more than 1,800 reproducing adults) in the lab of William Rice (University College Santa Barbara, USA). In 1995 (approximately 100 generations since establishment) the rearing protocol was changed to include non-overlapping generations, and a moderate rearing density with 16 adult pairs per vial (56 vials in total) during 2 days of adult competition, and 150-200 larvae during the larval competition stage^11^. In 2005, a copy of LH_M_ population sample was transferred to Uppsala University, Sweden (approximately 370 generations since establishment), and in 2012, to the University of Sussex (UK), when the current set of 223 haplotypes were sampled. At the point of sampling we estimate that the population had undergone 545 generations under laboratory conditions, 445 of which had been using the same rearing protocol.

Hemiclonal lines were established by mating groups of five clone-generator females (C(1)DX,*y*,f; T(2;3) *rdgC st in ri p^P^ bw*^D^) with 230 individual males sampled from the LH_M_ base population (see^1^). A single male from each cross was then mated again to a group of five clone-generator females in order to amplify the number of individuals harbouring the sampled haplotype. Seven lines failed to become established at this point. The remaining 223 lines were maintained in groups of up to sixteen stock hemiclonal males in two vials that were transferred to fresh vials each week. Stock hemiclonal males were replenished every six weeks by mating with groups of clone-generator females. A stock of reference genome flies (Bloomington *Drosophila* Stock Center no. 2057) was established and maintained initially using five rounds of of sib-sib matings before expansion. 223 virgin reference genome females were then collected and mated to a single male from each of 223 hemiclonal lines. Female offspring from this cross therefore have one copy of the reference genome and one copy of the hemiclonal haplotype. Groups of these hemiclonal females were collected as virgins, placed in 99% ethanol and stored at -20°C prior to DNA extraction.

### 2.2 DNA extraction

One virgin female per hemiclonal line, was homogenised with a microtube pestle, followed by 30-minute mild-shaking incubation in proteinase K. DNA was purified using the DNeasy Blood and Tissue Kit (Qiagen, Valencia, CA), according to manufacturer’s instructions. Volumes were scaled-down according to input material mass of input material. Barrier pipette tips were used throughout, in order to minimise cross-contamination of DNA. Template assessment using the Qubit BR assay (Thermo Fischer, NY, USA) indicated double-stranded DNA, 10.4Kb in length at concentrations of 2-4 ng/μl (total quantity 50-100ng).

### 2.3 Whole-genome resequencing

Sequencing was performed under contract by Exeter Sequencing service, University of Exeter, UK. The sonication protocol for shearing of the DNA was optimised for low concentrations to generate fragments 200-500bp in length. Libraries were prepared and indexed using the Nextera Library Prep Kit (Illumina, San Diego, USA). All samples were sequenced on a HiSeq 2500 (Illumina), with five individuals per lane. We also sequenced DNA from two individuals from the in-house reference line (Bloomington *Drosophila* Stock Centre no. 2057). One was prepared as the hemiclones, using the Illumina Nextera library (sample RGil), and the other using an older, Illumina Nextflex method (sample RGfi). The median number of read pairs across all samples was 29.23×10^6^ (IQR 14.07×10^6^). Quality metrics for the sequencing data were generated with FastQC v0.10.0 by Exeter Biosciences, and used to determine whether results were suitable for further analyses. For twelve samples with less than 8×10^6^ reads, sequencing was repeated successfully (H006, H041, H061, H084, H086, H087, H092, H098, H105), with a further three samples omitted entirely (H015, H016, H136), leaving 220 hemiclonal samples in total. As shown in Figures 1A and 1B, the read quality score and quality-per-base for the the samples taken forward for genotyping in this study were were well within acceptable standards, and similar across all samples.

**Figure 1.**
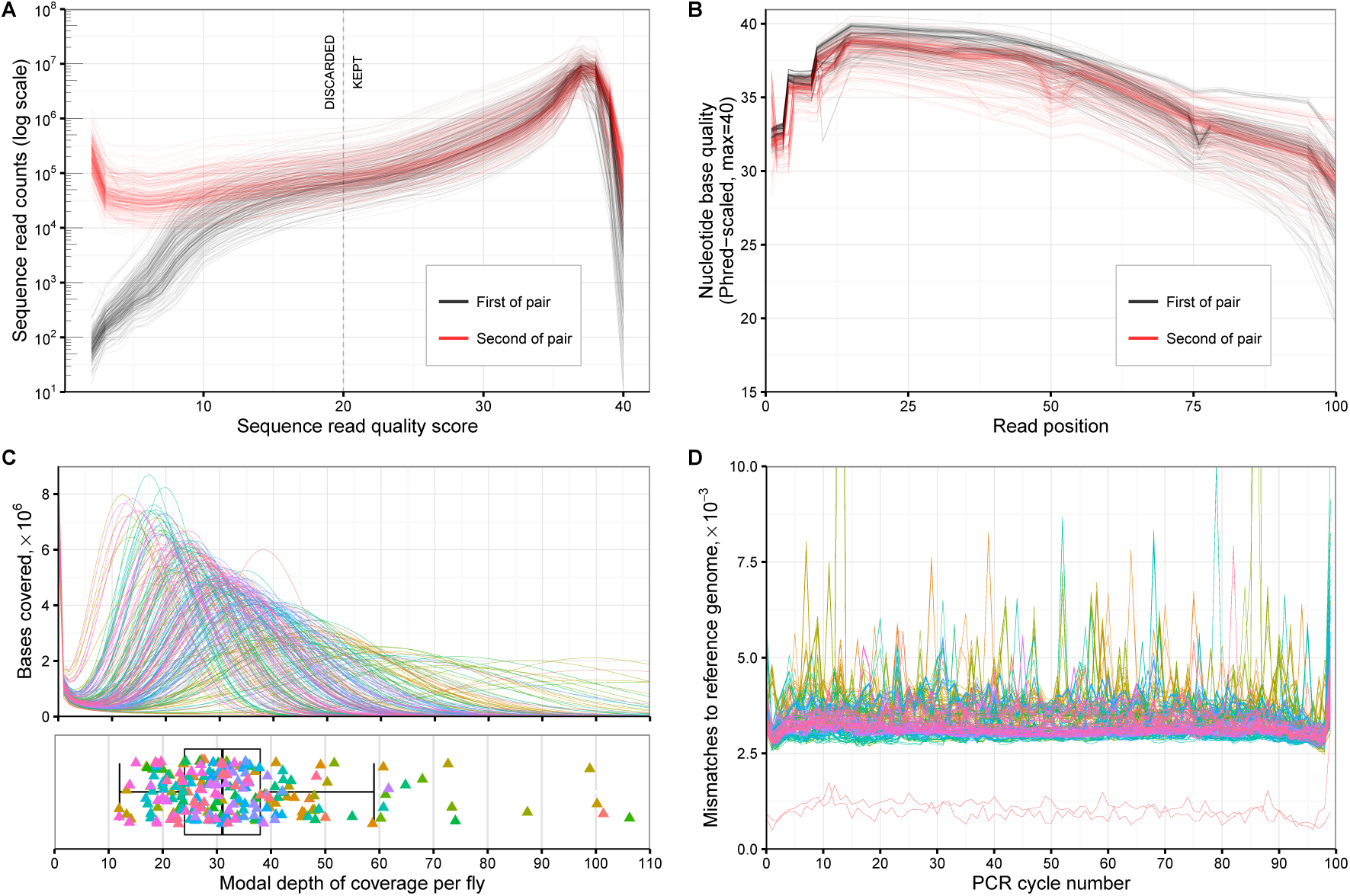
Next-generation sequencing assessment. A: Sequence read quality for each sample sequenced. Y-axis scale is logarithmic. B: Quality of sequences by nucleotide base position for each sample. C: Read depth of coverage distribution across each sample. Colouring corresponds to the order which which the samples were originally sequenced. D: Mis-matches to the dm6 reference genome assembly, by PCR cycle-number. Colouring is by sample as in plot C. The two red lines with visibly-lower mismatch rates than the others correspond to the two in-house BDGP/dm6 reference lines that were sequenced. Data and code for this figure is located at https://doi.org/10.5281/zenodo.159282.

### 2.4 Read mapping

Raw data *(fastq* files) were stored and processed in the Linux Sun Grid Engine in the High-Performance Computing facility, University of Sussex. Adaptor sequences (Illumina Nextera N501-H508 and N701-N712), poor quality reads (Phred score <7) and short reads were removed using Fastq-mcf (ea-utils v.1.1.2). Settings were: log-adapter minimum-length-match: 2.2, occurrence threshold before adapter clipping: 0.25, maximum adapter difference: 10%, minimum remaining length: 19, skew percentage-less-than causing cycle removal: 2, bad reads causing cycle removal: 20%, quality threshold causing base removal: 10, window-size for quality trimming:1, number of reads to use for sub-sampling: 3×10^5^.

Cleaned sequence reads were mapped to the *D. melanogaster* genome assembly, release 6.0 (Assembly Accession GCA_000001215.4^6^) using Burrows-Wheeler Aligner *mem* (version 0.7.7-r441)^9^, with a mapping quality score threshold of 20. Fine mapping was performed with both Stampy v1.0.24^10^ and the Genome Analysis Tool-Kit (GATK) v3.2.2^4^ (following^8^). Removal of duplicate reads, indexing and sorting was performed with Picard-Tools v1.77 and SamTools v1.0. The median depth of coverage across all samples used for genotyping was 31X (IQR 14, see Figure 1C). As shown in Figure 1D, the mean nucleotide mis-match rate to the dm6 reference assembly for the LH_M_ hemiclones was 3.27×10^−3^ per PCR cycle (IQR 0.2×10^−3^)contrasting with the two reference line samples for which the mis-match rate was 0.89 — 1.10×10^−^ per cycle. We observed spikes of nucleotide mis-matches in some PCR cycles for some samples, which are likely to be errors rather than true sequence variation.

### 2.5 Small-variant detection methods

Single-nucleotide polymorphisms (SNPs) and insertion/deletions (indels) ¡200bp in length, were detected and genotyped relative to the BDGP+ISO1/dm6 assembly, on chromosomes 2,3,4,X, and mitochondrial genome using Haplotyper Caller (GATK v3.4-0)^15^. Individual bam files were genotyped, omitting reads with a mapping quality under 20, stand call and emit confidence thresholds of 31, then combined and genotyped again. 143,726,002 bases of genomic sequence were analysed from which 1,996,556 variant loci were identified consisting of 1,581,341 SNPs, 196,582 deletions, and 218,633 insertions. Functional annotation was added using SNPeff^3^.

We used hard-filtering to remove variants generated by error, because the alternative, variant recalibration’ requires prior information on variant positions from a similar population or parents. Quality filtering thresholds were decided following inspection of the various sequencing metrics associated with each variant locus, and by software developers’ recommendations^15^. The filtering thresholds were: Quality-by-depth >2, strand bias (-log_10_.pFisher) <50, mapping quality >58, mapping quality rank sum >-7.0, read position rank sum >-5.0, combined read depth <15000, and call rate >90%. This filtering removed 167,319 variants (8.3%), leaving 1,829,237. Summary values for the variant quality metrics are shown in Table 1. Distributions of quality metrics for Haplotype Caller variants are shown in Supplementary Figure 2. The density of sequence variants, measured as the median for windows of 10Kb in length across the genome, was 75 per for biallelic SNPs, 1 for multi-allelic SNPs, 6 for biallelic indels, and 3 for multi-allelic indels (see Figure 2A). Mean separation between variants of any type or allele frequency was 78bp. As shown in Figure 2B the allele frequency distribution for biallelic SNPs and indels was similar, and broadly within expectations for an out-bred diploid population sample. The two in-house reference line individuals had 515 homozygous and 3171 heterozygous mutations from the reference assembly. The median genotype counts for the 220 LH_M_ hemiclone individuals, were 585 homozygous, 728,214 heterozygous and 4963 no-call (IQR 400, 36707 and 7876). Genotype counts for each individual are shown in Figure 2C.

**Table 1.** Haplotype Caller variant quality metrics and genotype frequencies.

| Variant type | SNPs (biallelic) | SNPs (multi) | Indels (biallelic) | Indels (multi) |
| --- | --- | --- | --- | --- |
| N | 1,411,395 | 43,798 | 138,687 | 65,660 |
| Total depth | 6440 (1725) | 6316 (2100) | 6134 (1836) | 5973 (2081) |
| Event length | 0 (0) | 0 (0) | 2 (5) | 1 (8) |
| Strand bias | 1.12 (2.25) | 1.34 (3.14) | 1.76 (3.88) | 1.77 (4.45) |
| Mapping quality | 62.12 (6.18) | 64.94 (8.57) | 71.17 (12.77) | 69.58 (11.36) |
| Map qual rank sum | 0.25 (1.04) | 0.9 (2.37) | 3.14 (3.21) | 2.68 (2.91) |
| Quality-by-depth | 16.65 (3.51) | 17(3.81) | 18.52 (6.21) | 16.96 (6.39) |
| Quality | 34968 (62236) | 57028 (67558) | 25842 (59889) | 40479 (63590) |
| Genotype counts |  |  |  |  |
| Reference | 151 (120) | 102(122) | 166(114) | 122(123) |
| Heterozygous | 70 (118) | 117(121) | 54(114) | 95(122) |
| Homozygous non-ref. | 0 (0) | 0(0) | 0(0) | 0(0) |
| No call | 0 (1) | 1(4) | 0(2) | 2(5) |

Values show the total number of variants, median (and IQR) for each metric. Data generated from *vcf* file using GATK VariantsToTable, on the quality-filtered data. Code and data used to generate this table located at https://doi.org/10.5281/zenodo.159282.

**Figure 2.**
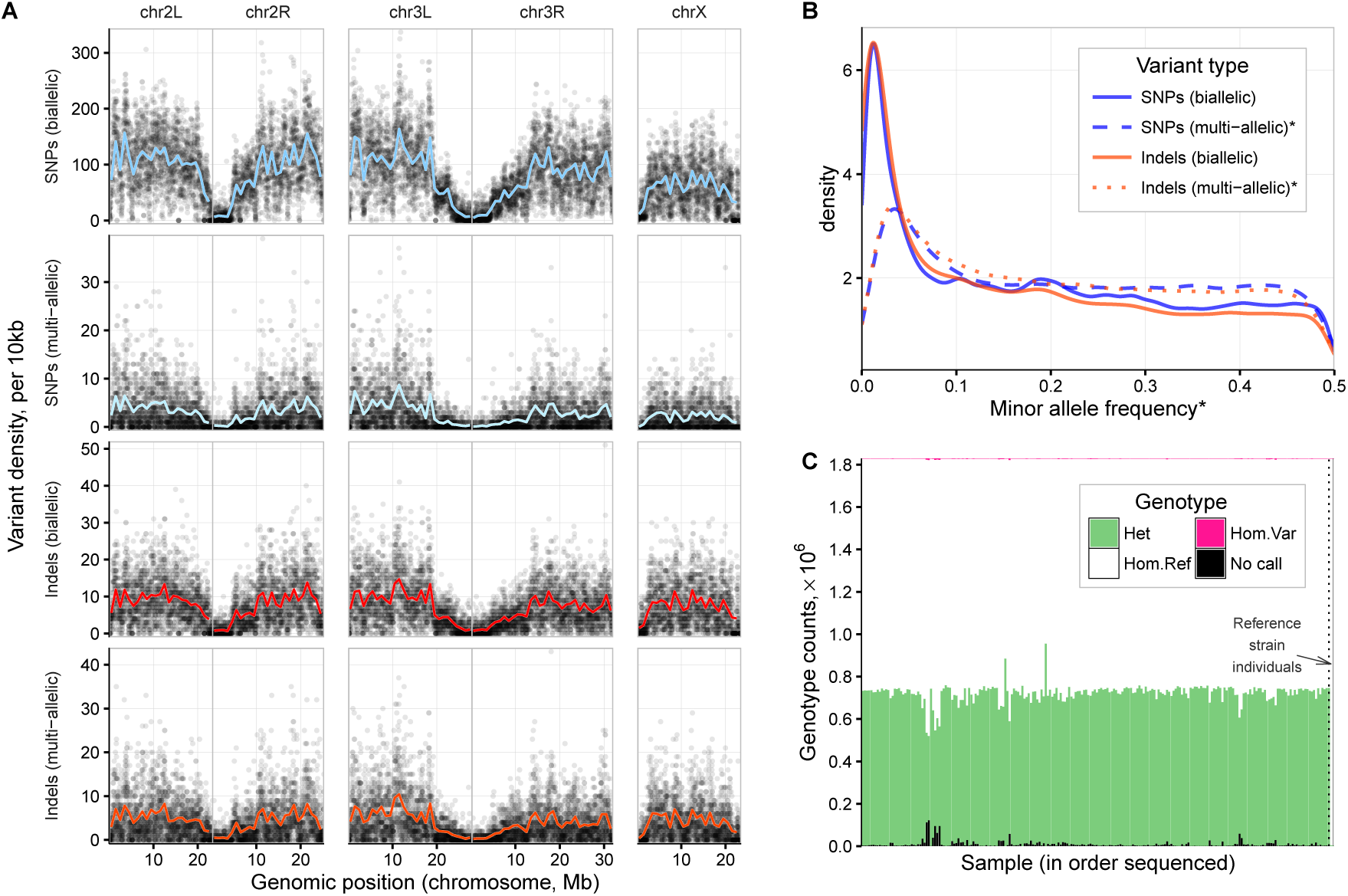
Summary of SNPs and indels in the LHM sample. A: Density of common variants across the genome (MAF>0.05 (Variants from the in-house reference line are included but account for less than 3,686 of the 1,825,917 common variants plotted (<0.2%). B: Allele frequency distribution by variant type. *MAF values were calculated from the count of heterozygous calls, and so for multi-allelic variants, the MAF is derived from the combined count of both alternate alleles. C: Genotype counts per individual genotyped. Data generated using GATK/3.4 VariantEvaluation function. Data and code for this figure is located at https://doi.org/10.5281/zenodo.159282.

For data submission to dbSNP, we removed 44,644 indels that were multi-allelic or greater than 50bp in length, and a further 57,662 variants that had null alternate alleles (likely due to being situated within a deletion). The genotype data submitted to dbSNP consists of 1,726,931 quality-filtered, functionally-annotated variant records (1,423,039 SNPs and 303,892 short, biallelic insertion and deletion variants) corresponding to 383,378,682 individual genotype calls.

### 2.6 Structural-variant detection methods

Large genomic variants – deletions and duplications, between 1Kb and 100Kb in length – were detected and genotyped using GenomeStrip v2.0^5^. One of the reference strain individuals (sample RGfi) was omitted from the this analysis because a different sequencing library preparation method was used to the other samples (see above). We included the following settings (according to developers’ guidelines): Sex-chromosome and k-mer masking when estimating sequencing depth, computation of GC-profiles and read counts, and reduced insert size distributions. Large variant discovery and genotyping was performed only on chromosomes 2, 3, 4 and X, omitting the mitochondrial genome and unmapped scaffolds.

We used the Genomestrip CNV Discovery pipeline with the settings: minimum refined length 500, tiling window size 1000, tiling window overlap 500, maximum reference gap length 1000, boundary precision 100, and genotyped the results with the GenerateHaploidGenotypes R script (genotype likelihood threshold 0.001). Following visualisation of the genotype results and comparison with the bam sequence alignment files using the Integrated Genomics Viewer (IGV)^13^, we excluded telomeric and centromeric regions where the sequencing coverage was fragmented, and six regions of multi-allelic gains of copy-number with dispersed break-points, previously reported to undergo mosaic *in vivo* amplification prior to oviposition^14^ (see Supplementary Table 1 for genomic positions, and Supplementary Figure 3 for visualisation of in vivo amplification in a sequence alignment file). We excluded 6 samples (H082, H083, H090, H097, H098, H153) for which 80-90% of the genome was reported by Genomestrip to contain structural variation, which we regarded as error. Most these samples were grouped by the order in which they processed for DNA extraction and sequencing, so this may have been caused partly by a batch-effect leading to differences in read pair separation, depth-of-coverage, and response to normal fluctuations GC-content.

Following removal of these samples, there were 2897 CNVs (1687 deletions, 877 duplications, and 333 of the ’mixed’ type), ranging in size from 1000bp to 217,707bp. We observed eight regions, for which Genomestrip identified multiple adjacent CNVs in single individuals, but which are likely single CNVs, 100Kb to 1.3Mb in length (Supplementary Table 2).

Using a combination of assumptions based on our breeding design, visualisation of read ’pile-ups’ across possible CNV regions using IGV, and inspection of quality metric distributions we used the following criteria for quality filtering: Quality score >15, Cluster separation <17, GC-fraction >0.33, no mixed types (deletions and duplications only), homozygous non-reference genotype count >0, heterozygous genotype count <200. Summaries of the quality metrics for quality-filtered data are shown in Table 2, and Supplementary Figure 2. We applied an upper limit to the cluster separation to remove groups of outliers in the upper end of the distribution, although this may have excluded many true, low-frequency variants. However, data on rare variants are not directly useful for our further investigations.

**Table 2.** Quality metrics for Genomestrip CNVs

| metric | Deletions | Duplications |
| --- | --- | --- |
| N | 78 | 89 |
| GC-fraction | 0.39 (0.07) | 0.42 (0.06) |
| Cluster separation | 8.84 (3.70) | 9.78 (3.17) |
| Quality | 103.93 (505.71) | 490.95 (1128.32) |
| Heterozygote count (max 213)* | 22.00 (42.50) | 42.00 (53.00) |
| Length (kb) | 2.20 (3.54) | 3.40 (2.35) |

Values show the total number of variants, median (and IQR) for each metric. Data generated from *vcf* file using GATK VariantsToTable, on the quality-filtered data. \*No CNVs in the quality-filtered samples had a 'no-call' or homozygous non-reference genotype.

After filtering, 167 CNVs remained (78 deletions and 89 duplications, size range 1Kb-26.6Kb). The positions and genotypes of these CNVs for each individual are shown in Figure 3. The genotype data for quality-filtered CNVs were combined with the data from 2252 indels >50bp from the Haplotype Caller pipeline, and a total of 2419 variants were uploaded to the public database on structural variation, NCBI dbVar. Although we have used methods for detecting SNPs, indels and CNVs, variants between 200bp and 1Kb are not reported by either HaplotypeCaller or Genomestrip. Additionally, sequence inversions are not detected by these methods, and the upper limits to CNV detection using Genomestrip, based on the parameters and results of this study are 100Kb-1Mb.

**Figure 3.**
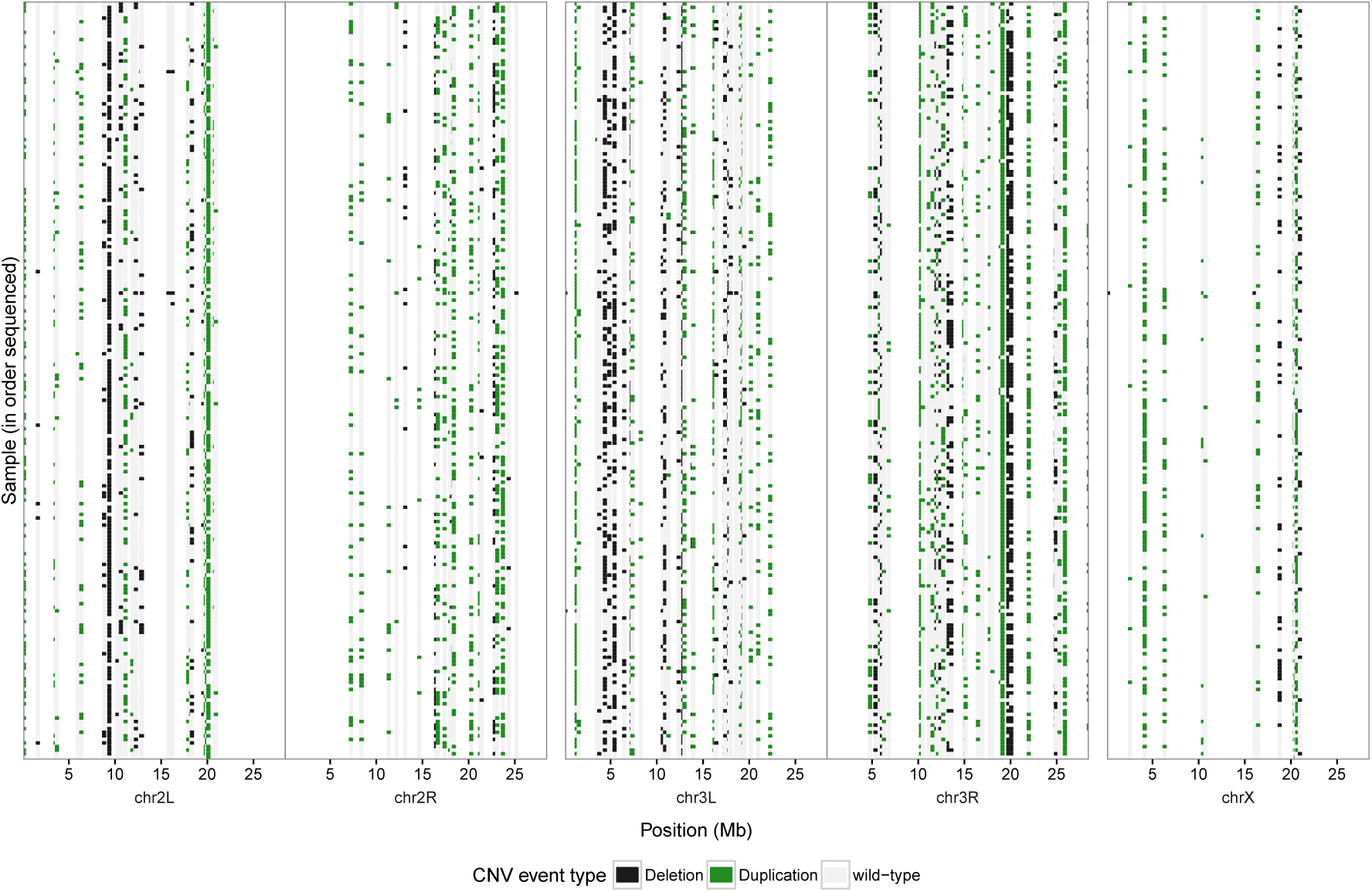
Structural variants across the *D. melanogaster* genome for the LHM population sample. Each row corresponds to an individual sequenced (in order originally sequenced from top to bottom, with the reference line at the bottom). Image generated using R/ggplot2 with data generated by GATK VariantsToTable with individual genotypes as copy-numbers. Data and code for this figure is located at https://doi.org/10.5281/zenodo.159282.

## 3 Dataset Validation

Initial validation of our methods can be seen by lack of variants in the two reference line individuals compared with the LH_M_ hemiclones (3,686 verses a median of 728,799 per sample). For a more thorough test the reproducibility of the genotyping and hemiclone method, we sequenced an additional hemiclone individual from three of the LH_M_ lines, and mapped the reads to the reference genome assembly as before. For HaplotypeCaller, we generated ’g.vcf’ files for each sample, and then performed genotyping and quality-filtering as described above, except that the original three samples were replaced with the replication test samples. Similarly, for Genomestrip, we performed structural variant discovery and genotyping all of the same samples as before, replacing three original samples with the replication test samples. We then used the GATK Genotype Concordance function to generate counts of genotype differences between the three pairs of samples. Overall results are presented in Table 3. Genotype reproducibility for quality-filtered bi-allelic SNPs was 98.5-99.5%, going down to 89.1-93.2% for filtered multi-allelic indels. Reproducibility of structural variant genotype calls was 95.6-100.0%, although we noted that for one individual (H119) filtering actually reduced the reproducibility rate from 99.7% to 95.6%. Full code, logs and numerical results can be found at http://doi.org/10.5281/zenodo.160539.

**Table 3.** Genotype reproducibility rates(%)*.

| Variant type | Sample ID | Unfiltered | Filtered |
| --- | --- | --- | --- |
| *HaplotypeCaller/3.4* |  |  |  |
| Bi-allelic SNP | H119 | 98.9 | 99.5 |
|  | H137 | 97.7 | 98.5 |
|  | H151 | 97.8 | 98.3 |
| Multi-allelic SNP | H119 | 95.0 | 96.6 |
|  | H137 | 92.3 | 94.0 |
|  | H151 | 92.1 | 93.6 |
| Bi-allelic indel | H119 | 98.1 | 98.6 |
|  | H137 | 96.3 | 96.8 |
|  | H151 | 96.0 | 96.4 |
| Multi-allelic indel | H119 | 91.9 | 93.2 |
|  | H137 | 88.0 | 89.3 |
|  | H151 | 87.9 | 89.1 |
| *Genomestrip/2.0* |  |  |  |
| Deletion | H119 | 99.7 | 95.6 |
|  | H137 | 100.0 | 100.0 |
|  | H151 | 100.0 | 100.0 |
| Duplication | H119 | 99.7 | 100.0 |
|  | H137 | 99.9 | 100.0 |
|  | H151 | 99.6 | 100.0 |

\*Presented values are the overall genotype concordance, as generated using GATK/3.4 Genotype Concordance function. Code, logs and output data are available at http://doi.org/10.5281/zenodo.160539.

Although these results indicate that our genotype accuracy is very good, there are several caveats to consider. In the quality-filtered small-variant data, seven samples (H034, H035, H040, H038, H039, H188, H174) had prominently higher genotype drop-out rates than the others (of 2-7%), as well as a higher proportion of homozygous non-reference genotypes (2-4%; See Figure 2C). Additionally two samples had prominently more heterozygous variants (H072:885,551 and H093:955,148 verses the other LH_M_ hemiclones: mean 710,934).

Although the genotype replication rate for the structural variants was also very high, we cannot exclude the possibility that, due to incomplete masking of hard-to-sequence regions of the reference assembly, variants which are artefacts reported in the original genotype data, may also be present in the replication genotype data.

## 4 Data Availability

All publicly-available records are for 220 LH_M_ hemiclone individuals and 2 in-house reference line individuals, with the exception of the large-variant data for which one in-house reference line sample and six LH_M_ hemiclones were omitted. The NCBI Bio-Project identifier is PRJNA282591. Code, logs and quality control data for each dataset, and for generating the figures and tables in this manuscript are publicly-available at the research data repository, Zenodo, https://zenodo.org/, ’Sussex Drosophila Sequencing’ community. Use of the files uploaded to Zenodo is under Creative Commons 4.0 license.

### 4.1 Data record 1: Sequencing data

Raw *fastq* sequence reads, and *bam* alignment files for the *D. melanogaster* are publicly-available at the NCBI Sequence Read Archive, accession number SRP058502. The code for read-mapping, alongside the run logs and quality-control data are available at https://doi.org/10.5281/zenodo.159251. Additionally the sequence alignment files for the corresponding *Wolbachia* have accession number SRP091004, with further supporting files at https://doi.org/10.5281/zenodo.159784.

### 4.2 Data record 2: Small-variant data

Records of quality-filtered sequence variants identified by GATK HaplotypeCaller in the LH_M_ hemiclones, and in the in-house reference line, have been submitted to NCBI dbSNP, https://www.ncbi.nlm.nih.gov/projects/SNP/snp_viewBatch.cgi?sbid=1062461, handle: MORROW_EBE_SUSSEX. In compliance with NCBI dbSNP criteria, variants >50bp in length, multi-allelic indels, and variants with a null alternate allele have been omitted. Genotype data, pre- and post-filtering, are also available at https://doi.org/10.5281/zenodo.159272, alongside the analysis code, run logs and quality-control data summaries.

### 4.3 Data record 3: Structural-variant data

Records of quality-filtered variants detected by GenomeStrip, and variants >50bp detected by Haplotype Caller are publicly-available at NCBI dbVar, accession number nstd134, http://www.ncbi.nlm.nih.gov/dbVar/nstd134. Unfiltered and filtered geno-type data, code for CNV discovery and genotyping using Genomestrip/2.0, run logs, and summary data are publicly-available at https://doi.org/10.5281/zenodo.159472.

## Author contributions

EM conceived and supervised the experiment. EM, TP, IF, MW and WG designed the experiment. TP and IF established and maintained the lines, and carried out the DNA extractions. WG analysed the sequencing and genotype data. WG and MW developed the read-mapping and variant-calling procedures. WG and EM wrote the manuscript.

## Acknowledgements

Sequencing was performed under contract by Exeter University, DNA Sequencing Service (UK), who also provided analysis advice, http://www.exeter.ac.uk/business/facilities/sequencing/. Crucial computational support was provided by Jeremy Maris at the Centre for High-Performance Computing, University of Sussex, http://www.sussex.ac.uk/its/services/research/highperformance. Bob Handsaker (Harvard Medical School, USA) provided analysis advice for use of Genomestrip for structural variant detection.

## 7 Supplementary information

### 7.1 URLs for External data and Software

dm6 Reference assembly (GCA_000001215.4) ftp://hgdownload.cse.ucsc.edu/goldenPath/dm6/

FastQC 0.10.0 http://www.bioinformatics.babraham.ac.uk/

EA-Utils (cleaning of sequence reads) 1.1.2 https://code.google.com/p/ea-utils/

Burrows-Wheeler Aligner (BWA) 0.7.7-r441 http://bio-bwa.sourceforge.net/

Stampy 1.0.24 http://www.well.ox.ac.uk/project-stampy

Genome Analysis Tool-Kit (GATK) 3.2.2, and later 3.4-0, as specified in the code and main manuscript text. https://www.broadinstitute.org/gatk/

PicardTools 1.77 http://picard.sourceforge.net

SamTools 1.0 http://samtools.sourceforge.net/

GenomeStrip 2.0 http://www.broadinstitute.org/software/genomestrip/

Script for generating genotype calls from GenomeStrip/2.0 CNV likelihood scores. More recent versions of Genomestrip include this script. ftp://ftp.broadinstitute.org/pub/svtoolkit/misc/cnvs/estimate_cnv_allele_frequencies.R

### 7.2 Supplementary Tables

**Table S1.** Regions from which structural variants reported by Genomestrip/2.0 were excluded.

| Chromosome | Start position | Stop position | Feature |
| --- | --- | --- | --- |
| 2L | 0 | 20,000 | telomere |
| 2L | 9,450,000 | 9,600,000 | *In vivo* amplification |
| 2L | 13,300,000 | 13,500,000 | *In vivo* amplification |
| 2L | 21,000,000 | 23,513,712 | centromere |
| 2R | 0 | 6,000,000 | centromere |
| 2R | 25,256,600 | 25,286,936 | telomere |
| 3L | 0 | 70,000 | telomere |
| 3L | 2,250,000 | 2,320,000 | *In vivo* amplification |
| 3L | 8,500,000 | 8,800,000 | *In vivo* amplification |
| 3L | 22,500,000 | 28,110,227 | centromere |
| 3R | 0 | 4,500,000 | centromere |
| 3R | 32,000,000 | 32,079,331 | telomere |
| X | 3,650,000 | 3,800,000 | *In vivo* amplification |
| X | 8,400,000 | 8,520,000 | *In vivo* amplification |
| X | 21,000,000 | 23,542,271 | centromere |

Genomic positions for centromeric and telomeric regions were determined following visualisation of *bam* sequence alignment files, where the sequencing coverage was fragmented, causing read pairs to be excessively separated without evidence of structural variation.

**Table S2.** Structural variants called as multiple events by Genomestrip

| Type | Chromosome | Start position* | Stop position* | Length(bp) | Sample present in |
| --- | --- | --- | --- | --- | --- |
| Duplication | 2L | 4,894,940 | 5,861,033 | 966,093 | H037 |
| Deletion | 2L | 15,335,536 | 16,655,783 | 1,320,247 | H023 |
| Deletion | 2R | 16,188,011 | 16,306,112 | 118,101 | H029 |
| Duplication | 2R | 21,499,905 | 22,386,557 | 886,652 | H165 |
| Deletion | 3R | 8,096,329 | 8,363,019 | 266,690 | H111 |
| Duplication | 3R | 15,720,028 | 17,043,150 | 1,323,122 | H148 |
| Duplication | 3R | 23,162,039 | 23,585,335 | 423,296 | H050 |
| Duplication | X | 19,995,505 | 20,112,715 | 117,210 | H203 |

\*Start and stop positions were determined from the limits of individual events identified by Genomestrip. Positions are relative to the *D.melanogaster* reference assembly dm6.

### 7.3 Supplementary Figures

**Figure S1.**
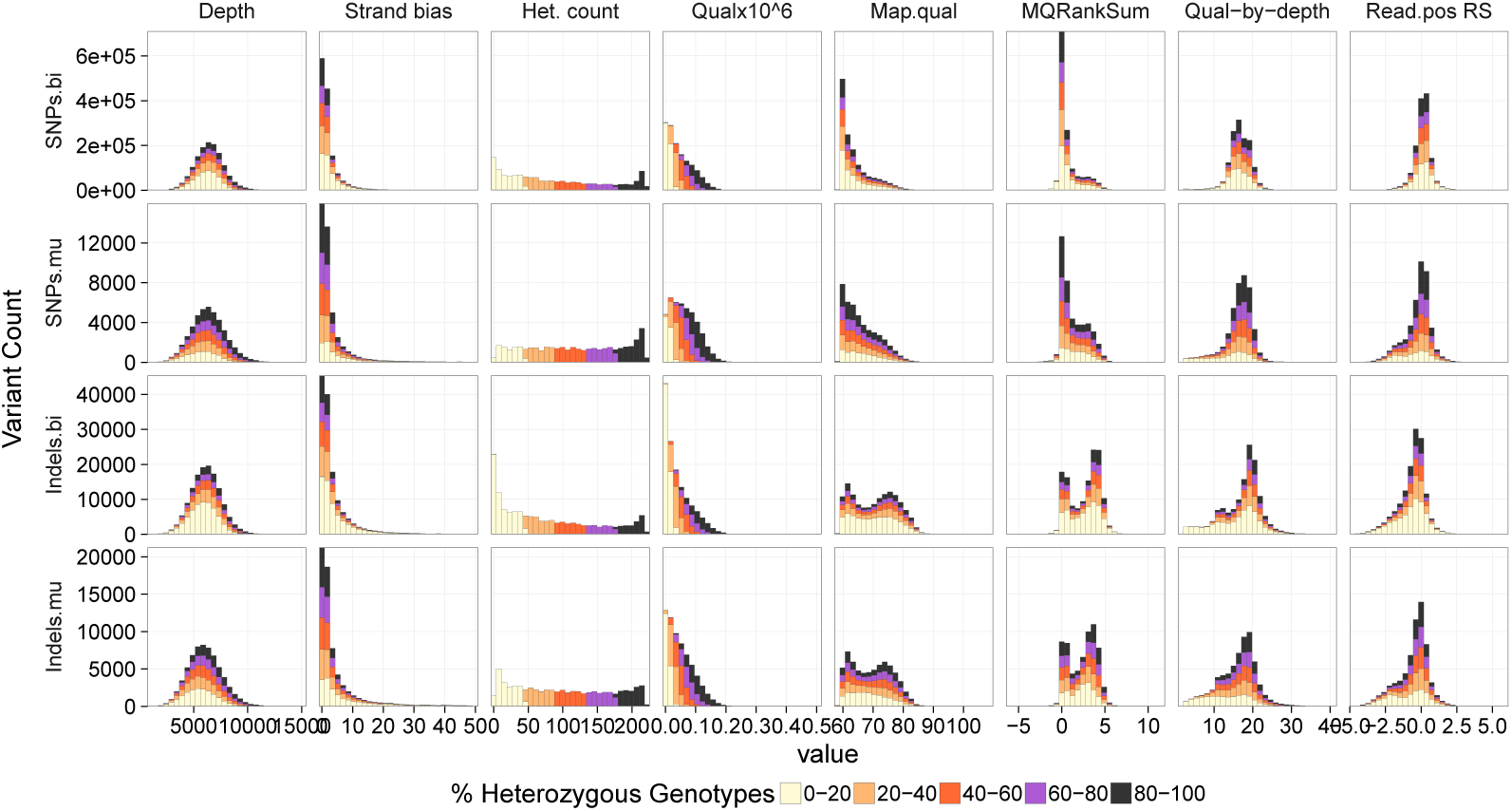
Distribution of quality metrics for SNPs and indels, detected by Haplotype Caller. Data generated by GATK VariantsToTable function and plotted in R. Plot bars are coloured by heterozygous genotype count, as a proxy for minor allele frequency in the hemiclone study sample. Code and data used to generate this figure are located at https://doi.org/10.5281/zenodo.159282.

**Figure S2.**
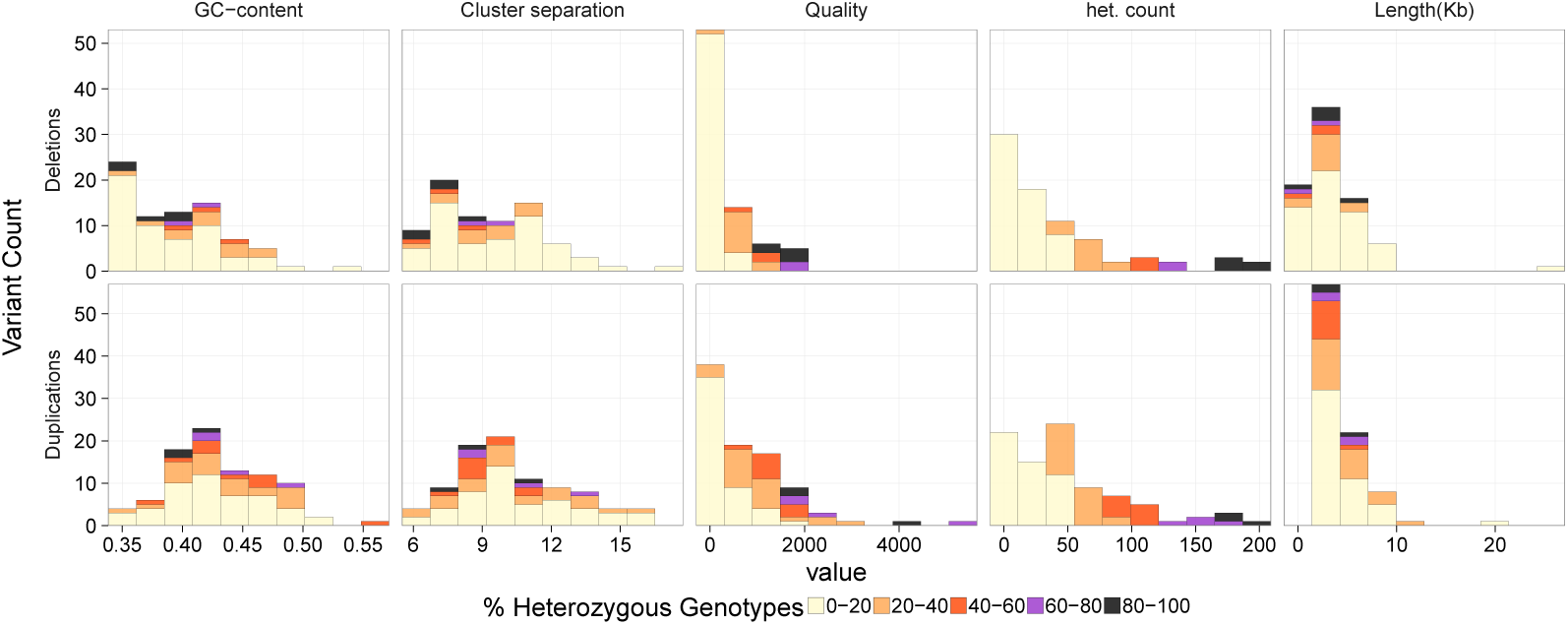
Distribution of quality metrics for structural variants detected by Genomestrip. Data generated by GATK VariantsToTable function and plotted in R. Plot bars are coloured by heterozygous genotype count, as a proxy for minor allele frequency in the hemiclone study sample. Data and code for this figure are located at https://doi.org/10.5281/zenodo.159282.

**Figure S3.**
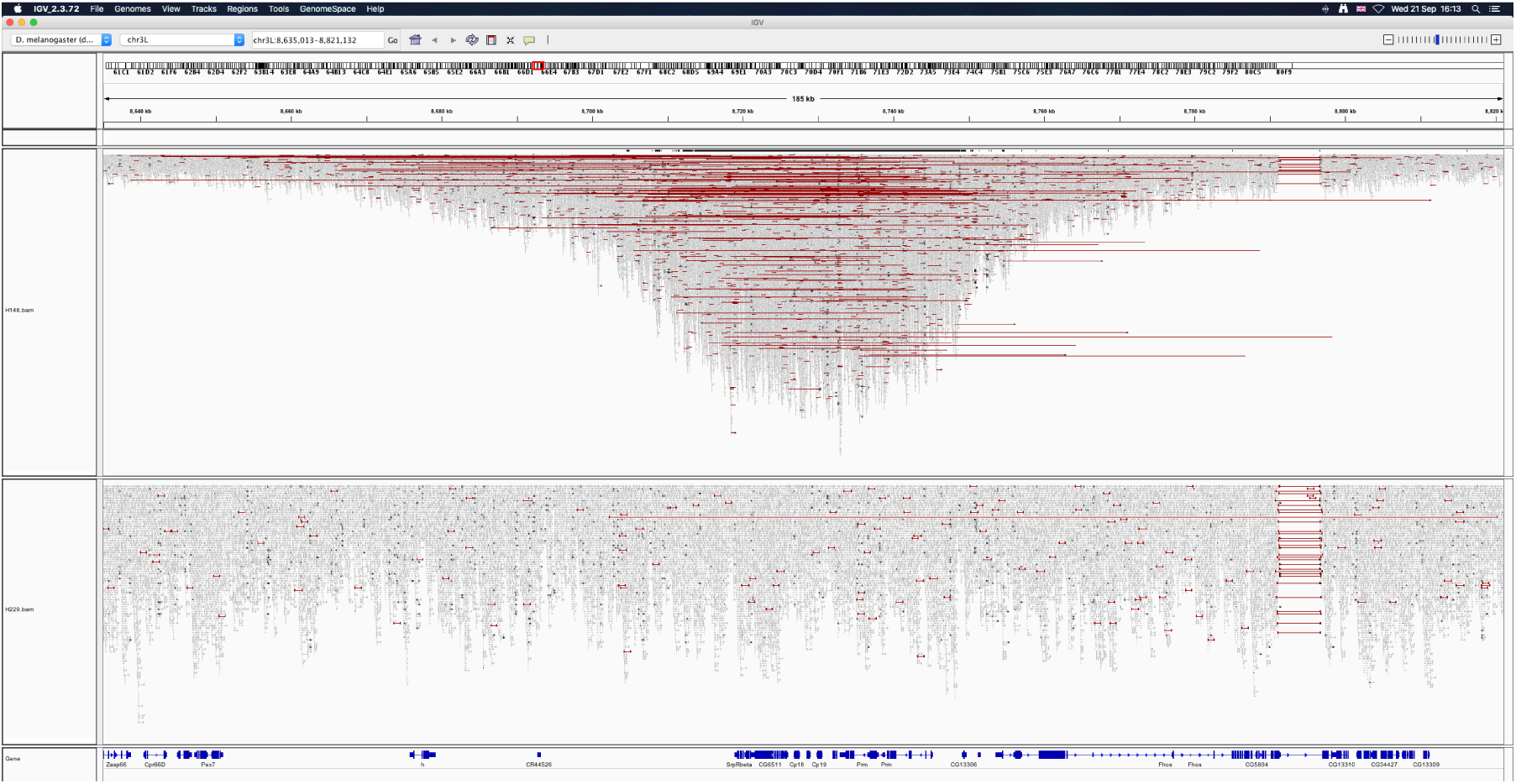
*In vivo* amplification in next-generation sequencing data. Image taken from visualisation of *bam* sequence alignment files using Integrated Genomics Viewer, and shows region around the chorion protein genes 18 and 19 on chromosome arm 3L.Small grey blocks indicate sequence reads. Horizontal red lines indicate read pairs which are >1000bp apart. The upper sample (H148) exhibits the amplification, whereas the lower sample (H001) does not. Also shown below in dark blue, are the positions of genes in the region.

